# Hexavalent Chromium Inhibits Nitrate-Dependent Anaerobic Methane Oxidation While Enriching Denitrifiers: Insights into Microbial Interactions for Simultaneous Methane, Nitrate and Chromate Removal

**DOI:** 10.1101/2025.06.17.660095

**Authors:** Yinxiao Ma, Garrett Smith, Suzanne S.C.M. Haaijer-Vroomen, Sanne Olde Olthof, Cornelia U. Welte, Martyna Glodowska

## Abstract

Chromate (Cr(VI)) is a widely distributed heavy metal known for its toxicity and adverse effects on human health. Due to its extensive industrial use, Cr(VI) is commonly detected in municipal and industrial wastewater, often co-occurring with other contaminants such as nitrate (NO_3_^−^). Nitrate-dependent anaerobic methane oxidation (N-DAMO) is a promising microbially driven process that enables the simultaneous removal of methane (CH_4_) and NO_3_^−^, potentially lowering the carbon footprint of wastewater treatment plants (WWTPs). Given that Cr(VI) can act as an alternative electron acceptor, its presence may influence the N-DAMO process either by supporting CH_4_ oxidation or, conversely, by exerting toxic effects that inhibit microbial activity. In this study, we investigated the impact of Cr(VI) on an N-DAMO enrichment culture composed of *Candidatus* Methanopereden*s* and *Candidatus* Methylomirabilis, using NO_3_^−^ as the electron acceptor and ^13^C-CH4 as the sole electron donor. Cultures were exposed to varying concentrations of Cr(VI), and microbial activity was assessed through GC-MS, 16S rRNA gene sequencing and qPCR. Here we found that while Cr(VI) was reduced within the culture, this process was not coupled to CH_4_ oxidation. In fact, CH_4_ oxidation was significantly inhibited, and the relative abundances of *Ca*. Methanoperedens and *Ca*. Methylomirabilis declined under increasing Cr(VI) concentrations. Cr(VI) reduction was likely mediated by the flanking microbial community via two possible pathways: (i) enzymatically, through nitrate reductase activity in denitrifying bacteria, and (ii) abiotically, through chemical reaction with nitrite (NO_2_^−^), an intermediate of denitrification. These findings highlight the potential of N-DAMO microbes to confer functional resilience in contaminated environments. However, the sensitivity of methanotrophic organisms to Cr(VI) underscores a limitation in applying this process to wastewater treatment systems containing heavy metals.

## 1. Introduction

Chromium (Cr) is a toxic heavy metal commonly found in industrial wastewater that threatens human and ecosystem health. Its hexavalent form, Cr(VI), is associated with dermatitis and neurotoxicity, as well as oxidative stress, DNA damage and carcinogenic effects at the cellular level (Mishra & Bharagava, 2016; Wise et al., 2022). Inhalation exposure to Cr(VI) has been classified as a Group 1 carcinogen by the International Agency for Research on Cancer (Welling et al., 2015). Beyond its impact on human health via occupational exposure, environmental Cr(VI) contamination adversely affects ecosystem functions due to its toxicity to plants and aquatic animals and ultimately affects human health through biomagnification in the food chain (Jobby et al., 2018; Tang et al., 2021). Chromium can exist in an oxidation state ranging from -2 to +6, with trivalent Cr(III) and hexavalent Cr(VI) being the most common and abundant (Bokare & Choi, 2011). Cr(III) and Cr(VI) have different biological roles and emission sources. Compared to Cr(VI), which is highly toxic, Cr(III) is generally insoluble, poorly absorbed by cells, and does not accumulate in living tissues (Levina & Lay, 2019; Levis & Majone, 1981). Moreover, Cr(III) is usually absorbed by soil colloids and immobilized in organic matter and metal (hydr)oxides, hindering its migration in groundwater and natural environments (Yang et al., 2021). Therefore, reducing Cr(VI) to Cr(III) effectively limits chromium contamination. Cr(III) is ubiquitous in various natural waters and predominantly originated from the weathering of Cr-bearing minerals, such as chromite (FeCr_2_O_4_) and bentorite (Ca_6_(Cr, Al)_2_(SO_4_)3) (Ukhurebor et al., 2021). In contrast, anthropogenic activities such as stainless-steel production and the mining industry are the primary sources of Cr(VI)(Ramli et al., 2023). It was shown that stainless-steel smelting slags, a by-product of steel production, contain up to 10% Cr(VI), which can enter groundwater from landfills through infiltration (Gu et al., 2021; Spooren et al., 2016). Recent reports from a Dutch regional supervisory authority have revealed Cr(VI) in groundwater at a depth of five meters beneath a steel plant in the Netherlands (Kreling & Schoorl, 2022). The mining industry is also an important source of Cr(VI) pollution (Ramli et al., 2023). Examples from the mining region in Sukinda, India, have demonstrated that mining operations can cause significant water contamination, with Cr(VI) concentrations reaching 2.48 mg/L in surface water and 1.35 mg/L in groundwater (Chowdhury et al., 2016). Due to its broad applications in various branches of industries, Cr(VI) is among the most common heavy metals found in the environment (Ramli et al., 2023). Consequently, many countries have regulations on Cr discharge, for example, the EU has implemented uniform emission standards for Cr(VI) and total Cr are 1 mg/L and 5 mg/L, respectively (Tumolo et al., 2020; Vaiopoulou & Gikas, 2020).

Nitrate-dependent anaerobic methane oxidation (N-DAMO) is a process that couples the reduction of NO_3_^−^ to dinitrogen gas (N_2_) and the oxidation of methane (CH_4_) to carbon dioxide (CO_2_), enabling the complete removal of NO_3_^−^ in methanogenic anoxic wastewater (Ettwig et al., 2010). In combination with other microbial treatments, N-DAMO was proposed as a more sustainable alternative for wastewater treatment plants (WWTPs) (Fan et al., 2020). Current research on N-DAMO is mainly focused on optimizing the nitrogen removal efficiency and demonstrating the performance of N-DAMO under real wastewater scenarios with the co-occurrence of different pollutants (van Kessel et al., 2018; Wissink et al., 2024a). However, the effect of Cr(VI) on the efficiency of the N-DAMO process, and specifically on NO_3_^−^ and CH_4_ removal is still unknown.

On the one hand, strong oxidative stress and cellular toxicity may reduce or even completely inhibit nitrogen and methane removal via N-DAMO. A recent incubation study demonstrated that Cr(VI) of > 30mg/L altered the composition of heterotrophic denitrification inoculum and rapidly inhibited the NO_3_^−^ reduction (Wang et al., 2023), which might also be true for N-DAMO. However, the N-DAMO community consists mainly of methanotrophic archaea such as *Candidatus* Methanoperedens and bacteria such as *Candidatus* Methylomirabilis oxyfera and therefore, its response to Cr(VI) remains unknown. On the other hand, the N-DAMO community demonstrated remarkable resilience against lead contamination (Wissink et al., 2024) and has a genetic potential for a versatile range of electron acceptors for CH_4_ oxidation, including manganese (IV), arsenic (V), vanadium (V) and chromium (VI) (Glodowska et al., 2022a). Moreover, CH_4_ oxidation releases more energy when coupled with Cr(VI) reduction than NO_3_^−^ reduction to NO_2_^−^ under chemical standard conditions, which indicates Cr(VI) might be a thermodynamically more favorable electron acceptor for *Ca*. Methanoperedens (Eq.1 and 2) (Bhattarai et al., 2019).

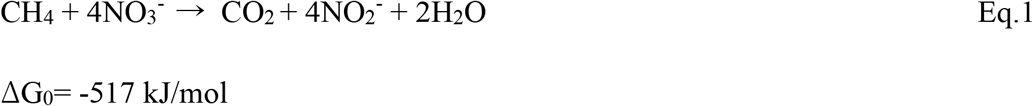

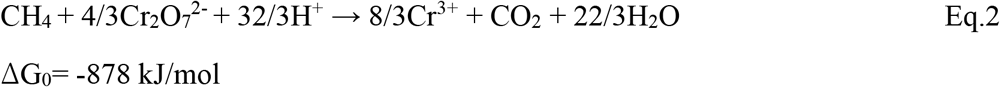

There is an ongoing debate on the role of Cr(VI) in CH_4_ oxidation. Al Hasin et al. were the first to report a simultaneous Cr(VI) reduction and CH_4_ oxidation by the pure culture of *Methylococcus capsulatus* under aerobic conditions (Al Hasin et al., 2010). In recent years some evidence suggested that Cr(VI) can be utilized as the sole electron acceptor by anaerobic methanotrophs such as *Candidatus* Methanoperedens. However, previous experiments, although conducted with anaerobic methanotrophic enrichment, did not directly link Cr(VI) reduction with conversion of CH_4_ to CO_2_, or Cr(VI) was not the only electron acceptor in the system (Lu et al., 2016; Luo et al., 2019). Only recently, it was demonstrated that indeed *Ca*. Methanoperedens can couple Cr(VI) reduction while oxidizing ^13^C-CH_4_ to ^13^C-CO_2_ (Wang et al., 2024).

Therefore, we performed a batch incubation experiment with an N-DAMO enrichment culture to explore the potential of N-DAMO application further and assess the possibility of simultaneous NO_3_^−^ and Cr(VI) removal. Unlike in the previous studies, we applied two electron acceptors concurrently. We challenged the N-DAMO culture with different concentrations of Cr(VI) to investigate 1) the effect of different concentrations of Cr(VI) on the NO_3_^−^ reduction and CH_4_ oxidation rate and the composition of the N-DAMO community; 2) whether the N-DAMO community can use Cr(VI) as an alternative electron acceptor to oxidize CH_4_.

## 2. Materials and Methods

### 2.1 Batch incubation

A batch incubation experiment was conducted in 120 mL sterile glass serum bottles in biological triplicate to explore the response of the N-DAMO community to Cr(VI) and the potential of anaerobic CH_4_ oxidation coupled with Cr(VI) reduction. The batch incubation experiment was set up in an anoxic glovebox (97% N_2_ and 3% H_2_, O_2_ < 15 ppm). First, all bottles received 50ml of medium as described in (Kurth et al., 2019), 3mM (final concentration) of sodium nitrate (NaNO_3_), and 0.2 ± 0.004 g (dry weight) of N-DAMO inoculum. After that, potassium chromate (K_2_Cr_2_O_7_) solution was added to bottles to reach a 0.3, 0.7, and 1 mM final concentration of Cr(VI). Treatment without added K_2_Cr_2_O_7_ served as a control. Stock solutions of NaNO_3_, and K_2_Cr_2_O_7_ were gassed with N_2_/CO_2_ to remove dissolved O_2_ before use. All incubation bottles were closed with butyl rubber stoppers and aluminum crimp caps before being transferred from the glove box. The headspace gas of each incubation bottle was exchanged with a mixture gas of N_2_/CO_2_ (9:1 vol:vol), and finally, 0.4 mmol of ^13^C-CH_4_ was injected into the headspace of each bottle. The pressure in the incubation bottles exceeded two standard atmospheres (> 2 bar) at the beginning of the experiment to ensure the dissolution of CH_4_ in the liquid and to maintain anoxic conditions. Bottles were kept in the dark at room temperature for 263 hours. The N-DAMO community used in this study was first obtained from an agricultural ditch in The Netherlands, and after a long-term enrichment in a bioreactor, it was dominated by *Ca*. Methylomirabilis (∼26%) and *Ca*. Methanoperedens nitroreducens (∼44%) at the time of the experiment (Raghoebarsing et al., 2006; Schoelmerich et al., 2022).

### 2.2 Gas Analysis

At each timepoint, 20 µl gas samples were withdrawn in duplicate from the headspace of each bottle for CH_4_ and CO_2_ analysis. The concentration of ^13^C-CO_2_ and ^12^CO_2_ was measured by gas chromatography coupled to mass spectrometry (Trace DSQ II, Thermo Finnigan, Austin TX, USA), and headspace CH_4_ concentration was quantified by gas chromatography with flame ionization detection (Hewlett Packard HP 5890 SeriesII Gas Chromatograph, Agilent Technologies, California, USA). The total ^13^C-CO_2_ concentration was calculated using equation S1 (Supplementary Materials).

### 2.4 Liquid Phase Analysis

NO_3_^−^, NO_2_^−^, and dissolved Cr concentrations in the liquid phase of each bottle were monitored throughout the experiment. At each time point, sample collection was performed in the glovebox, and 0.5ml of the liquid sample was withdrawn with a sterile syringe and needle for NO_3_^−^ quantification with the Griess assay (Sun et al., 2003). Another 0.5ml of the liquid sample was mixed with 9.5mL of 1% HNO_3_ for Cr quantification by ICP-MS (8900, Agilent Technologies, USA). Because of the high solubility of Cr(VI) and the low solubility of Cr(III), in this study, the concentration of dissolved Cr was used as a proxy for the concentration of Cr(VI).

### 2.4 DNA extraction and microbial community analysis

At the end of incubation (263 hours), the sealed incubation bottles were opened in the glove box and shaken gently. Then, 2 ml of the biomass was transferred into an Eppendorf tube for following DNA isolation. The DNA extraction was performed using the PowerSoil DNA extraction kit (DNeasy PowerSoil Pro Kit, QIAGEN, Hilden, Germany) from 0.5 g of wet biomass following the manufacturer’s protocol. The DNA concentration was measured by Qubit® 2.0 Fluorometer with DNA HS kits (Life Technologies, Carlsbad, CA, USA). Only the DNA samples with a concentration higher than 20 ng/µL were used for the following analysis. 16S rRNA gene amplicon sequencing was performed by Macrogen (Amsterdam, The Netherlands) using the Illumina MiSeq Next Generation Sequencing platform. Paired-end libraries were prepared with the Illumina Herculase II Fusion DNA Polymerase and Nextera XT Index Kit V2 (Illumina, Eindhoven, Netherlands). Primers used for bacterial and archaeal 16S rRNA gene amplification are listed in Table 1.

**Table 1.**
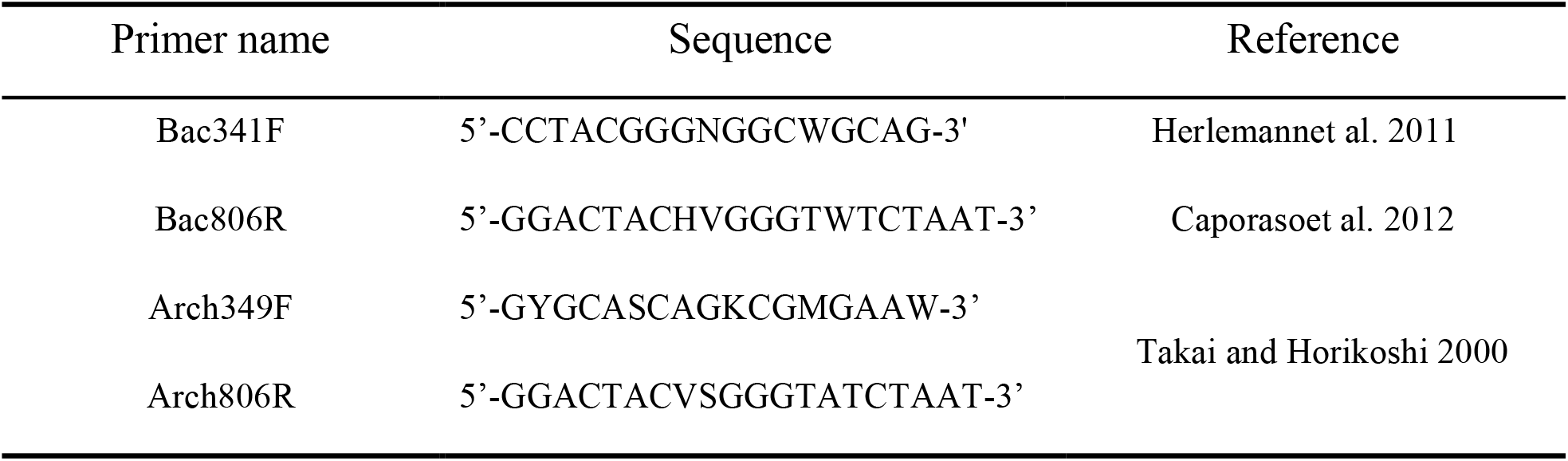
Bacterial and archaeal primers sequence.

For bacteria, original sequencing results were quality-filtered and trimmed to remove chimeric sequences (settings: left trim at 17 and 20, truncation length at 267 and 270, maxE 2), followed by denoising and dereplication (settings: error learning with 1e10 bases, pooling during denoising, and trimming overhangs during merging). Amplicon Sequence Variant (ASV) identification and read was then conducted, with taxonomic assignment performed using the SILVA version nr138 training set (Quast et al. 2013) and read abundance counting using DADA2 and its utilities (v1.22.0 (Callahan et al.2016) in R (v4.1.2; R Core Team, 2019).

### 2.5 Quantitative PCR

In the N-DAMO enrichment, following via 16S rRNA amplicon sequencing, *Ca*. Methanoperedens nitroreducens was found to be the only archaeal species present. Therefore, qPCR was used to track its abundance. To determine *Ca*. Methanoperedens 16S rRNA gene copy numbers, qPCR was performed using archaea 16S rRNA gene dsDNA gBlocks (Integrated DNA Technologies, USA) as standards for calibration and primers specific for *Ca*. Methanoperedens (Table 2) using a CFX96TM Real-Time System (C1000 TouchTM Thermal Cycler, Bio-Rad). A single qPCR reaction consisted of 5 µL 2x PerfeCTa SYBR Green FastMix (Quanta Bio), 800 nM of each primer, 2.4 µL Invitrogen™ Nuclease-Free Water (Thermo Scientific), and 1 µL of 0.1 ng DNA extracted from 0.5 g of wet biomass as templet. To generate a standard curve, a 16S archaea gBlock (Integrated DNA Technologies) was serial diluted in 10-fold steps Invitrogen™ Nuclease-Free Water (Thermo Scientific) resulting in a standard curve with concentrations of 1 ng/µL to 0.000001 ng/µL DNA. Each qPCR assay was performed in technical triplicate. The 16S rRNA gene qPCR program started with a single heating step to 98°C for 3 minutes followed by 40 cycles of 98°C for 10 seconds, 59°C for 15 seconds, and 72°C for 20 seconds. The PCR program ended with a melting curve generated ranging from 59°C to 98°C increasing with 0.5°C for 5 seconds each. A pmoA gene qPCR, was executed like the 16S RNA gene qPCR, using a pmoA gene dsDNA gBlock to generate a standard curve and using 400 nM of each pmoA gene primer (Table 2) in the reaction mix. The PCR program started with a heating step of 98 °C for 5 minutes, followed by 40 cycles of 98 °C for 10 seconds, 55 °C for 15 seconds, and 72 °C for 20 seconds. The PCR program ended with a melting curve ranging from 55 °C to 98 °C. For data analysis, CFX Maestro software v1.1 (Bio-Rad) and Excel (Microsoft) were used to calculate the gene copy number per 1 g of wet biomass.

**Table 2.**
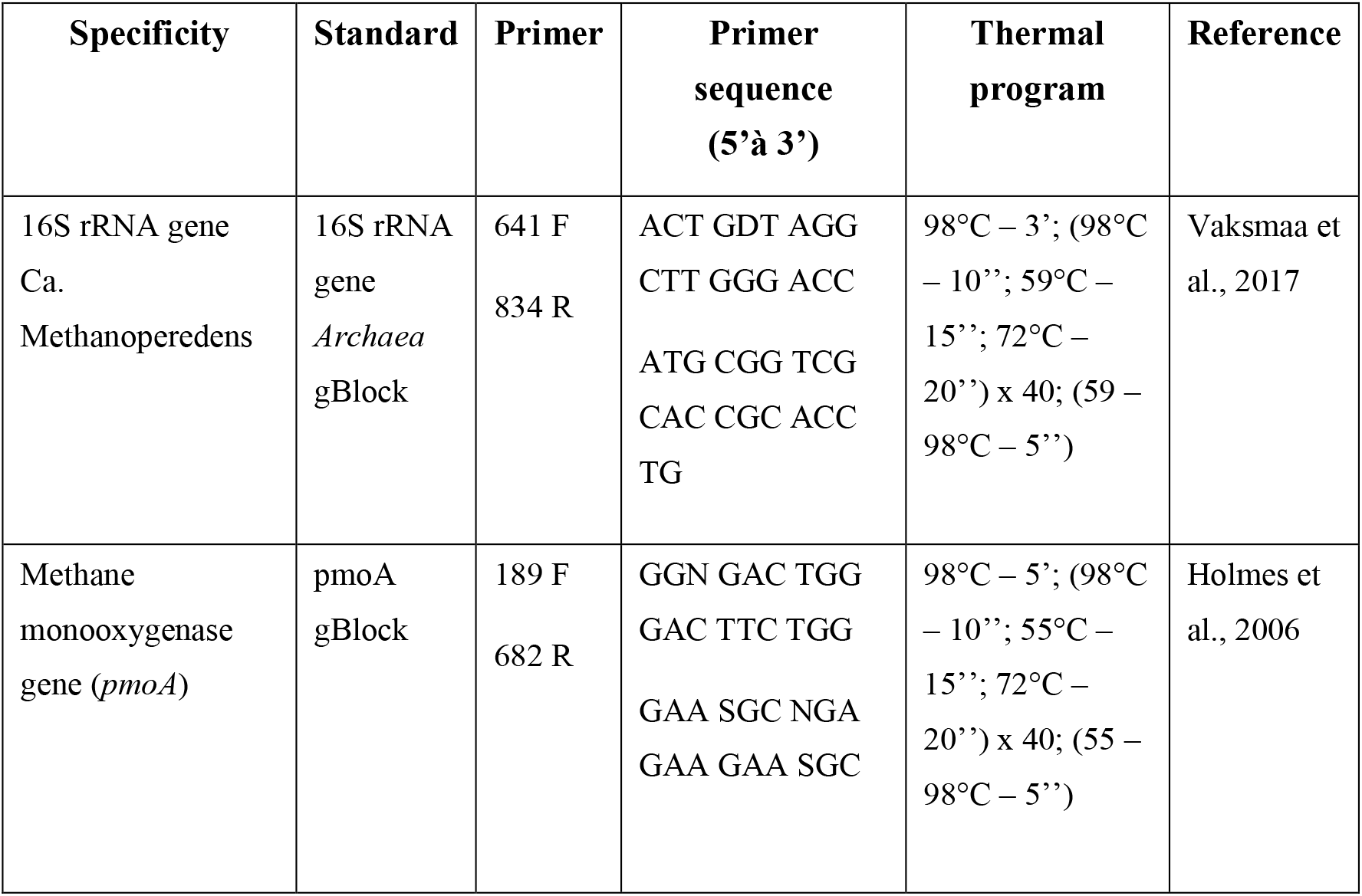
Primers, primer sequences and thermal programs used for quantification of *Ca*. Methanoperedens 16S rRNA gene copy numbers.

Differences in number of 16S rRNA Ca. *Methanoperedens* and *pmoA* gene copy among treatments with varying Cr(VI) concentrations were assessed using one-way analysis of variance (ANOVA), followed by Tukey’s Honestly Significant Difference (HSD) post-hoc test to identify pairwise differences between groups. The assumptions of normality and homogeneity of variances were verified prior to analysis. A significance level of α = 0.05 was used as the threshold to determine statistically significant differences.

## 3. Results and Discussion

### 3.1 Cr(VI) inhibits anaerobic CH_4_ oxidation

Cr(VI) inhibits anaerobic CH_4_ oxidation, which was determined by decreased CH_4_ consumption and lack of ^13^C-CO_2_ formation (Fig.1a and b). In the control setup, the N-DAMO enrichment culture was incubated with NO_3_^−^ as the sole electron acceptor and ^13^C-CH_4_ as the sole electron donor. In the first 80 hours, the headspace CH_4_ concentration decreased from 0.4 to 0.3 mM (Fig.1a) while ^13^C-CO_2_ increased from 0 to 60 µM (Fig. 1b). Simultaneously, the NO_3_^−^ was depleted from the initial 2.43 ± 0.55 mM, and no NO2^−^ accumulation was detected at the end of incubation (Fig.2a and b). Therefore, the stoichiometry of NO_3_^−^ reduction coupled with CH_4_ oxidation was at a ratio of 1.458 (2.43 mM*0.06L NO_3_^−^: 0.01 mM CH_4_), which is close to the theoretical stoichiometry of 1.6 of the overall reaction (Eq.5) of the stepwise NO_3_^−^ reduction to N_2_ by *Ca*. Methanoperedens (Eq.1) and *Ca*. Methylomirabilis (Eq.3) (Yao et al., 2024)

**Figure 1.**
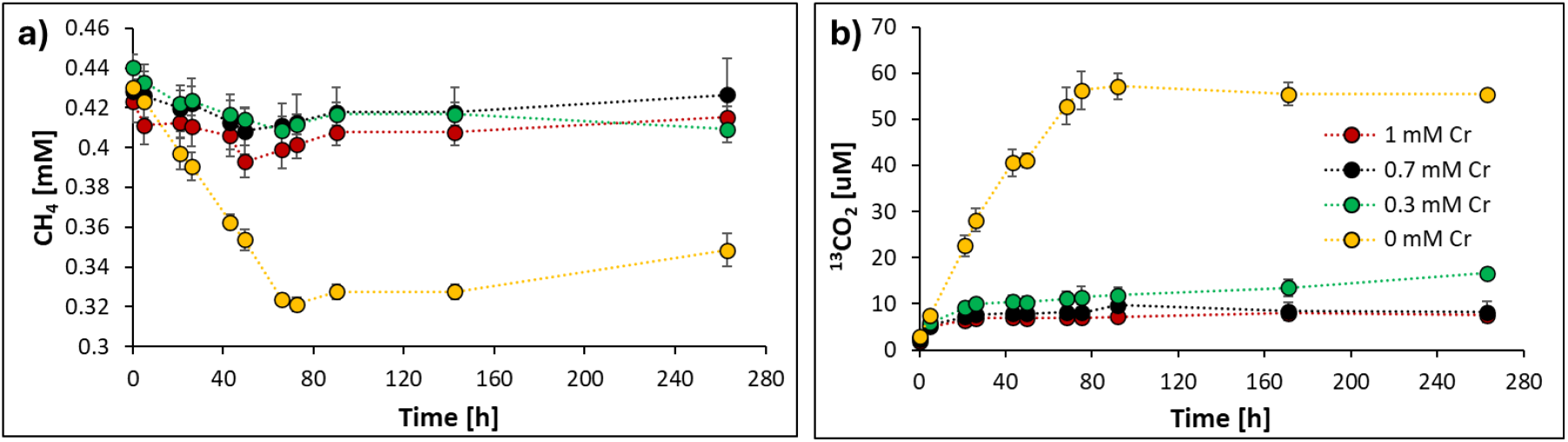
Changes of a) headspace CH_4_ concentration and b) total ^13^CO_2_ concentration under different Cr(VI) concentrations: 0 mM (yellow), 0.3 mM (green), 0.7 mM (black), and 1 mM (red). N-DAMO cultures were amended with ^13^CH_4_ and incubated under anoxic conditions at 30 °C. Each data point represents the mean ± standard deviation (SD) from three biological replicates (n = 3).

**Figure 2.**
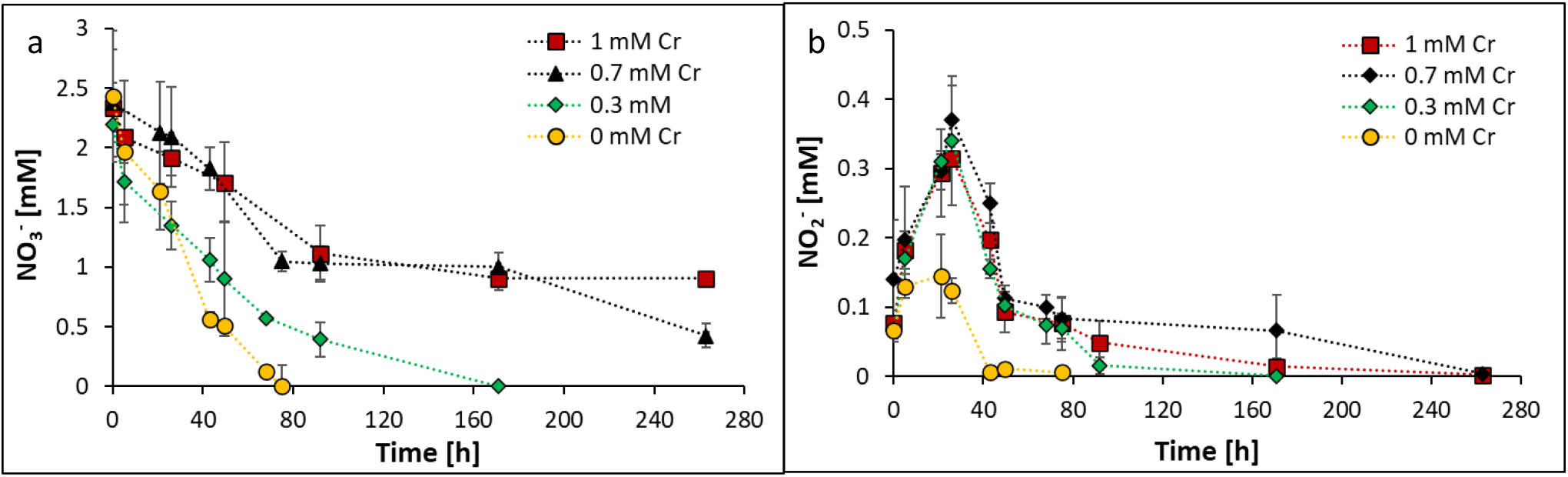
Changes in a) NO_3_^−^ and b) NO2^−^ concentration under different Cr(VI) concentrations: 0 mM (yellow), 0.3 mM (green), 0.7 mM (black), and 1 mM (red). Each data point represents the mean ± standard deviation (SD) from three biological replicates (n = 3).

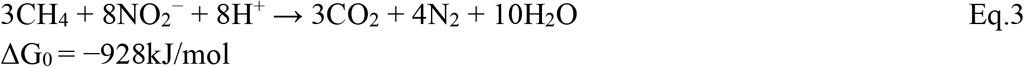

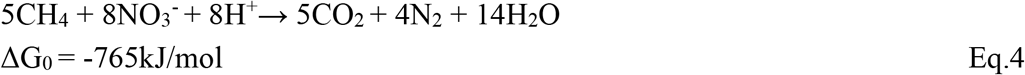

Moreover, the CH_4_ oxidation and ^13^C-CO_2_ production immediately stopped when NO_3_^−^ and NO_2_^−^ were depleted at 75h, confirming that the CH_4_ oxidation was coupled with NO_3_^−^ reduction, and an active N-DAMO process occurred (Fig.1a and 2a). As the N-DAMO community consisted of two main methanotrophs, *Ca*. Methanoperedens and *Ca*. Methylomirabilis, we assumed these taxa were key drivers of CH_4_ oxidation.

Only 60 μM of ^13^C-CO_2_ was produced during CH_4_ oxidation with only NO_3_^−^, suggesting that about 55% of consumed CH_4_ was converted into ^13^C-CO_2_ after 72 h (Fig.1a and b). This is presumably due to incomplete oxidation of CH_4_ by *Ca*. Methylomirabilis. This most abundant methanotrophic bacterium in our N-DAMO enrichment culture may exhibit incomplete CH_4_ oxidation under NO_2_^−^ limitation (Wu et al., 2015) producing methanol and other intermediate carbon compounds that can cross-feed the flanking community stimulating denitrification and Cr(VI) reduction.

*Ca*. Methanoperedens appears to be genetically equipped to use Cr(VI) as electron acceptor as it encodes for enzymes known to be involved in in bacterial reduction of Cr(VI) such as nitroreductases (Thatoi et al., 2014) or chromate reductase (Glodowska et al., 2022). Moreover, several previous studies suggested that Cr(VI) reduction can be coupled to CH_4_ oxidation, with some studies specifically pointing toward *Ca*. Methanoperedens as a key player in this process (Dong et al., 2019; Liu et al., 2025; S. Wang et al., 2024). However, only one recent study by (S.Wang et al., 2024) using isotope tracer experiment, electron microscopy, fluorescent visualization, and proteomic analysis provided strong evidence of *Ca*. Methanoperedens mediating this process independently from the flanking community. Contrary to previous expectations, the process was found not to involve chromate or nitrate reductases. Instead, numerous cytochrome *c* proteins were among the most upregulated, suggesting that extracellular Cr(VI) reduction occurs via multiheme cytochrome *c* (MHCs).

In our experiment CH_4_ oxidation significantly decreased (0.3 mM Cr) or was entirely inhibited (0.7, 1mM Cr) in the presence of Cr(VI). At the 0.3mM Cr treatment, only a small fraction of added ^13^C-CH_4_ (∼8%) was consumed, which is much lower than the 25% ^13^C-CH_4_ decrease in the control setup (Fig.1b). Specifically, with 0.3mM Cr(VI), only a slight decrease in CH_4_ concentration after 143h was observed, and the final amount of ^13^C-CO_2_ was only 30% of that when no Cr(VI) was added (Fig.1b). The higher concentration of Cr(VI) completely stopped the CH_4_ oxidation. No CH_4_ consumption nor ^13^C-CO_2_ production was observed in 0.7 and 1mM Cr(VI) treatments, apart from the initial decrease of headspace CH_4_ concentration caused by the dissolution of CH_4_ from the headspace to the liquid phase (Fig.1a). Although in our experiment Cr(VI) was decreasing over time (Fig. 3) this process was not coupled to CH_4_ oxidation.

**Figure 3.**
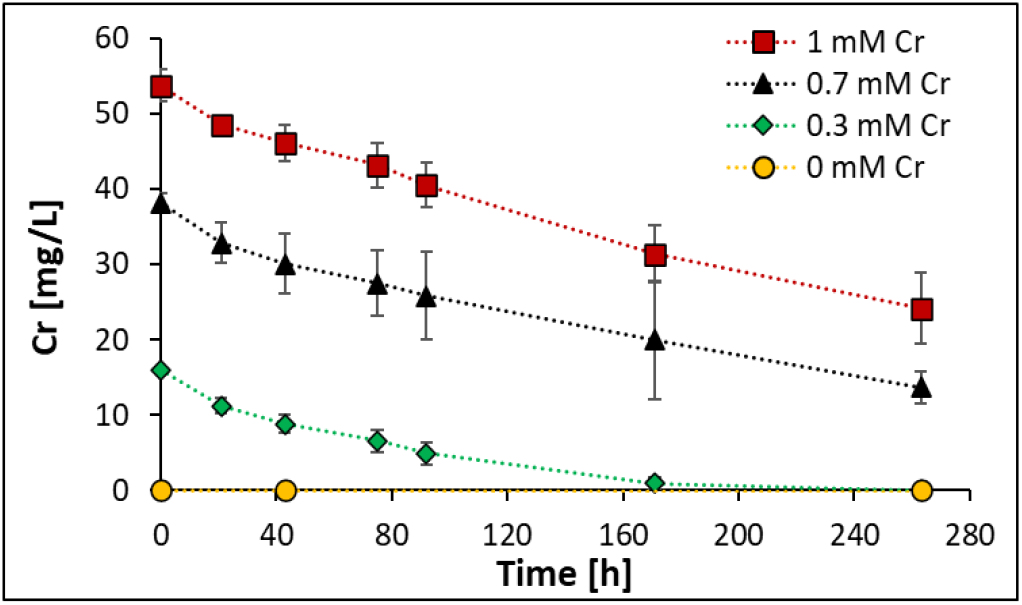
Changes in concentration of dissolved Cr over time: 0 mM (yellow), 0.3 mM (green), 0.7 mM (black), and 1 mM (red). N-DAMO cultures were amended with ^13^CH_4_ and incubated under anoxic conditions at 30 °C. Each data point represents the mean ± standard deviation (SD) from three biological replicates (n = 3). As most of the Cr(III) generated by microbial reduction is insoluble, the dissolved Cr in this experiment represents the Cr(VI) concentration (S. Wang et al., 2024).

This inhibitory effect of Cr(VI) on CH_4_ oxidation might be explained by the fact that in our study we used both electron acceptors (Cr(VI) and NO_3_^−^) simultaneously. Although chromate is thermodynamically a more favorable electron acceptor (Eq.2), the N-DAMO enrichment culture used in our experiment was continuously grown on NO_3_^−^ for many years, therefore most likely became better adapted to use it as electron acceptor. Considering that our experiment took only 263h (∼11 days), it was probably insufficient for the culture to adapt and switch to chromate as electron acceptor. In the 0.3 mM Cr treatment, after 120 h of incubation, the N-DAMO enrichment gradually restored its CH_4_ oxidizing capacity. By this time, Cr(VI) was nearly completely consumed, while NO_3_^−^ was still available (Fig. 2a and 3). It is, however, likely that after depletion of NO_3_^−^ and continuous supply of Cr(VI), eventually the process of CH_4_ oxidation would be coupled with Cr(VI) reduction.

Overall, it is evident that the presence of Cr(VI) at low concentration hinders and at high concentration, completely prevent CH_4_ oxidation even when NO_3_^−^ is available.

### 3.2 Nitrate reduction is hindered by Cr(VI)

Denitrification showed a much higher resilience against Cr(VI) toxicity compared to CH_4_ oxidation that was completely inhibited at higher Cr(VI) concentrations. In the control setup where no Cr(VI) was added, NO_3_^−^ was depleted within 75h and it was clearly coupled to CH_4_ oxidation (Fig.1). However, in the presence of 0.3mM Cr(VI), it took more than twice this time (171h) to remove all NO_3_^−^. In the 0.7 and 1 mM Cr(VI) treatments, NO_3_^−^ reduction was incomplete after the 263 h incubation period, with final concentrations of 0.43 ± 0.10 mM and 0.91 ± 0.10 mM, respectively (Fig. 2a). This clearly shows that NO_3_^−^ reduction was mediated by more diverse and less susceptible flanking community rather than just N-DAMO. Furthermore, the Cr(VI) amendment also caused a higher accumulation of NO_2_^−^. In the presence of Cr(VI), almost two-fold higher NO_2_^−^ concentrations (∼0.35mM) were measured compared to the control (∼0.15mM) at 26h (Fig.2b). However, this accumulation of NO_2_^−^ appeared to be transient and independent of Cr(VI) concentration as there was no difference in the NO_2_^−^ concentration (∼0.35mM) between the three Cr(VI) concentrations, and in all treatments, NO_2_^−^ concentrations were below the detection limit at the end of incubation (Fig.2b).

Denitrifiers are known to have a higher tolerance to toxic Cr(VI), and many studies have demonstrated their ability to reduce a wide range of heavy metal oxides, including Cr(VI) (Sahinkaya et al., 2017; Zhou et al., 2021). The extracellular polymeric substance (EPS) secreted by some denitrifiers can form a protective layer to slow down Cr(VI) penetrating the cell membrane (Aquino & Stuckey, 2004). Besides, the versatile enzymes of denitrifiers such as nitrate reductases, nitrite reductase, and flavoproteins may transfer electrons to Cr(VI), facilitating the reduction of Cr(VI) to less toxic Cr(III) (Zhou et al., 2021). Previous studies have shown that Cr(VI) concentrations of about 0.4 mM typically do not adversely affect denitrification. Moreover, many microorganisms retain over 80% of their denitrification capacity even at higher Cr(VI) concentrations ranging from 0.95–1.5 mM (Jin et al., 2017; Q. Wang et al., 2023). This is likely a case in our experiment as well, where NO_3_^−^ reduction decreased with increasing concentration of Cr(VI). Nevertheless, even at the highest Cr concentration more than 60% of NO_3_^−^ was consumed. We assume that decrease in NO_3_^−^ reduction was due to adverse effects of Cr(VI) on the methanotrophic denitrifiers rather than heterotrophic denitrifiers in the flanking community which were less affected by the presence of Cr(VI).

### 3.3 Cr(VI) reduction is mediated by N-DAMO flanking community

The total Cr concentration in the solution was measured to investigate whether the N-DAMO enrichment 1) can cope with Cr(VI) toxicity and 2) has a metabolic potential to use it as electron acceptor. As most of the reduced Cr(III) should precipitate from the solution or form insoluble complexes with organic matter (S. Wang et al., 2024; Zhou et al., 2021), we assume that total Cr measured in the solution is equivalent to the Cr(VI) concentration. A steady decrease in Cr(VI) concentration was observed in all treatments (Fig.3) despite the methanotrophic activity being largely or entirely inhibited by the Cr(VI), evidencing that Cr(VI)-reduction was not directly linked to CH_4_ oxidation. *Ca*. Methanoperedens and other methanotrophs have previously been shown to synthesize carbon storage compounds, such as polyhydroxyalkanoates (PHAs). Notably, *Ca*. Methanoperedens has demonstrated the ability to generate electric current using PHAs, suggesting that these compounds can serve as electron donors for the reduction of alternative electron acceptors (X. Zhang et al., 2022). Therefore, it is possible that a small portion of Cr(VI) was reduced by methanotrophs using PHAs, rather than CH_4_, as the electron source.

All Cr amendment treatments demonstrate a nearly identical Cr(VI) reduction rate of about 0.12-0.14 mg*L^-1^*h^-1^ in the first 92 hours independently of the starting concentration of Cr. This further indicates the strong tolerance and high Cr(VI) reduction efficiency of the flanking community in the N-DAMO enrichment under the high level of Cr(VI) (Fig.3).

Methane was the only electron donor used in our experiment, nevertheless NO_3_^−^ and Cr(VI)-reduction were clearly fueled by another electron donor. We hypothesize that organic carbon necessary to power the heterotrophic community originated either from dead biomass (necromass) or intermediate carbon compounds such as acetate were produced via partial CH_4_ oxidation under rate-limiting conditions (Cai et al., 2019).

### 3.4 Cr(VI) altered microbial community structure

To investigate changes in microbial community composition in the presence of Cr(VI), DNA was extracted at the end of the incubation and bacteria and archaea 16S rRNA amplicon sequencing was performed (Fig. 4). Additionally, to get a deeper insight on the response of N-DAMO methanotrophs a qPCR assays were carried out (Fig. 5). Since *Ca*. Methanoperedens was the only archaeal taxon detected in our experiment, *Ca*. Methanoperedens 16S rRNA gene was used as a marker. Gene encoding for particulate methane monooxygenase (*pmoA*) was used as a proxy for the abundance of *Ca*. Methylomirabilis.

**Figure 4.**
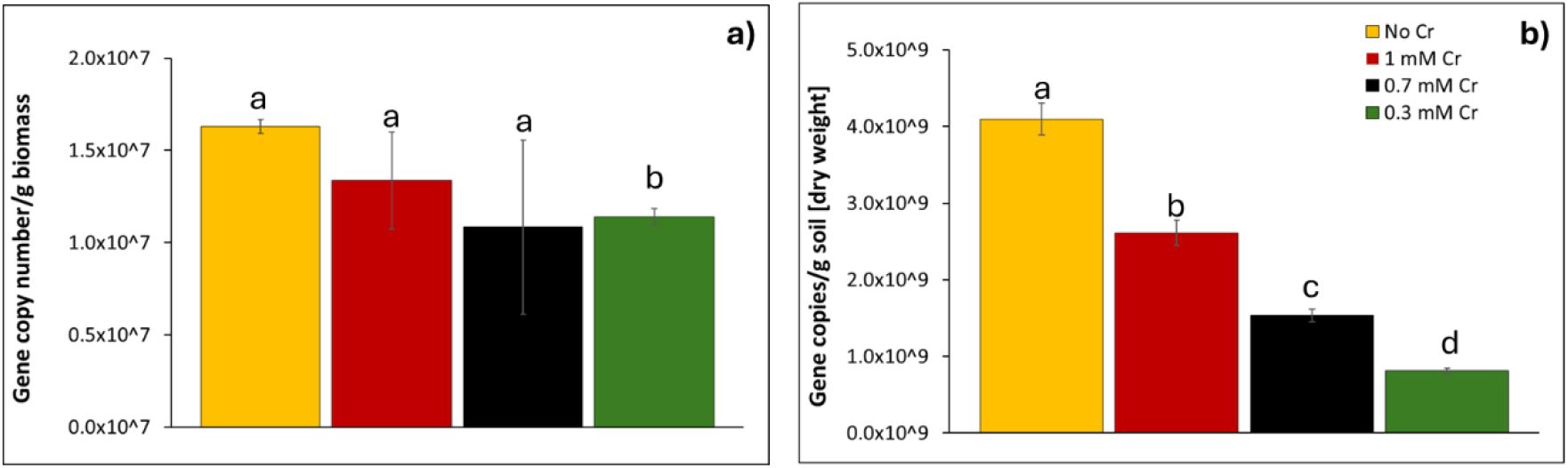
Effect of Cr(VI) on a) *Ca*. Methanoperedens 16S rRNA and b) *pmoA* gene abundance expressed in gene copy number/g of wet biomass. Different letters were assigned to groups where the means are significantly different (*p* < 0.05). Groups that share a letter are not significantly different from each other. Error bars represent standard deviation from three measurements.

**Figure 5.**
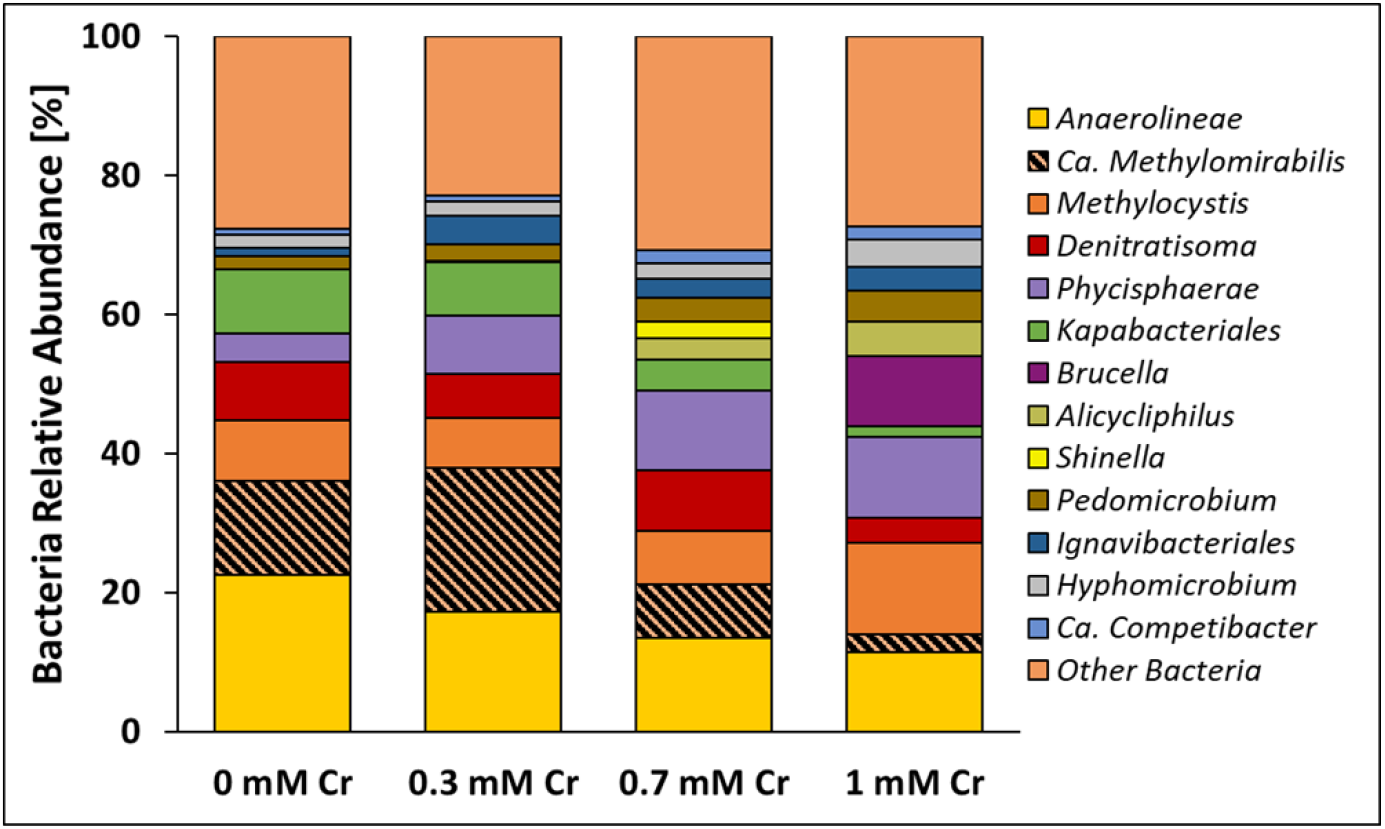
Bacterial community composition under different Cr(VI) concentrations. Stacked bar chart showing the relative abundance of bacterial taxa based on 16S rRNA gene amplicon sequencing from N-DAMO enrichment cultures incubated with 0, 0.3, 0.7, and 1 mM Cr(VI). DNA samples were collected at the end of the incubation period (273 h).

The results showed that Cr(VI) exposure negatively affected *Ca. Methanoperedens*, as indicated by reduced gene copy numbers compared to the control (Fig. 4a). Although all Cr(VI) treatments led to a decrease in gene abundance, the most substantial reduction was observed at 0.3 mM Cr(VI), which was significantly lower than the control (*p* < 0.005). Interestingly, the highest gene copy number among the Cr(VI)-supplemented cultures was found in the 1 mM treatment, suggesting a non-linear response to Cr(VI) concentration.

Cr(VI) also had a negative impact on *pmoA* gene copy numbers which were significantly lower (*p* < 0.005) in all Cr(VI) treatments compared to the control, implying negative effect of Cr(VI) on *Ca*. Methylomirabilis. Similar to *Ca. Methanoperedens*, the lowest *pmoA* gene copy number was detected at 0.3 mM Cr(VI), and the highest at 1 mM, again indicating a non-dose-dependent effect.

These results are consistent with 16S rRNA amplicon sequencing data (Fig. 5), where the relative abundance of *Ca*. Methylomirabilis initially increased at 0.3 mM Cr(VI) (21%) compared to the control (13.5%) but declined sharply to 7.7% and 2.6% in the 0.7 and 1 mM Cr(VI) treatments, respectively. It is however important to mention that *Ca*. Methylomirabilis was not the only taxon in our microbial community encoding for *pmoA* gene. The *pmoA* gene is also found in *Methylocystis*, a Type II methanotroph (Knief, 2015), which in our experiment appeared unaffected by Cr(VI). In fact, its relative abundance increased to 13% in the 1 mM Cr(VI) treatment compared to 8% in the control. Therefore, the elevated *pmoA* gene copy number at 1 mM Cr(VI) is likely attributable to the increased abundance of *Methylocystis* rather than *Ca. Methylomirabilis*.

Overall, the two anaerobic methanotrophs, *Ca*. Methanoperedens and *Ca*. Methylomirabilis appeared to be negatively affected by Cr(VI) presence. This decreased abundance of methanotrophs was also reflected in the lack of anaerobic CH_4_ oxidation particularly visible in 0.7 and 1mM Cr treatment.

In addition, Cr(VI) also altered the abundance of flanking microbial community in N-DAMO enrichment. Previously mentioned, *Methylocystis*, an aerobic methanotroph, showed the highest enrichment (13%) in the 1 mM Cr treatments implying its resilience to toxic effect of Cr(VI) and potential involvement in Cr(VI) reduction, as oxygen was not present in the incubation bottles. A recent study by (Q. Liu et al., 2025) demonstrated that *Methylocystis* can mediate NO_3_^−^ and Cr(VI) reduction in collaboration with denitrifying bacteria to support CH_4_ oxidation under micro-aerobic conditions. Similarly, taxa related to *Phycisphaerae* belonging to *Planctomycetota* phylum increased its abundance in all Cr treatments reaching 11.6 % in 1mM Cr(VI) concentration compared to the control where it represented only 4% microbial community. On the other hand, *Kapabacteriales* accounting for 9% of microbial communities in the control decreased their abundance to 7.6, 4.5 and 1.6% in 0.3, 0.7 and 1mM Cr treatment, respectively, likely due to its vulnerability to Cr(VI) toxicity. Heterotrophic denitrifier *Denitratisoma* previously suggested to be able to reduce heavy metal oxides (Lu et al., 2022), remained relatively stable, reaching its maximum abundance of 8.6% in the 0.7 mM treatment. Another denitrifier, *Alicycliphilus*, was undetectable in the control but progressively increased in abundance from 0.3, 3 to 5% with increasing concentration of Cr treatment. We suspect that this microorganism was involved in Cr(VI) reduction, particularly as *Alicycliphilus* was previously reported to be able to transform Cr(VI) to Cr(III) and was prevalent in polluted sites such as landfills and wastewater sludges (Lyu et al., 2021; Solís-González & Loza-Tavera, 2019).

Several taxa showed increased abundance with rising concentrations of Cr(VI), suggesting their potential involvement in Cr(VI) reduction. Notably, this included the iron- and manganese-oxidizing genus *Pedomicrobium* (Braun et al., 2009), members of the order *Ignavibacteriales*, and *Hyphomicrobium*, which were previously observed to be abundant in a methanotrophic reactor supplied with NO_3_^−^ and Cr(VI) as electron acceptors (Q. Liu et al., 2025). Additionally, denitrifying bacterium *Ca*. Competibacter was also enriched under these conditions.

### 3.5 Competitive inhibition of denitrification by Cr(VI)

Our batch incubation results indicate that Cr(VI) - a potential electron acceptor and a toxic heavy metal - exerts differential inhibitory effects on N-DAMO and denitrifying microbial communities. Cr(VI) exhibits acute toxicity toward the N-DAMO process, causing an inhibition of CH_4_ oxidation and decrease in relative abundance of anaerobic methanotrophs. In contrast, its effect on denitrifiers was different, aligning more closely with the characteristics of competitive inhibition (Almeida et al., 1995; W. Zhang et al., 2014).

Previously, some studies have proclaimed that Cr(VI)/Cr(III) is a more favorable electron acceptor than NO_3_^−^/NO_2_^−^, due to its significantly higher redox potential under standard chemical conditions. (ΔE^0^_NO3−/NO2−_: +0.83V vs ΔE^0^Cr(VI)/Cr(III) :+1.33V) (Ng et al., 2022; Patrick & Jugsujinda, 1992). Therefore, when Cr(VI) and NO_3_^−^ coexist in the system, Cr(VI) should be preferentially reduced. This is also supported by previous observations that NO_3_^−^ reduction is slower in the presence of Cr(VI), whereas the Cr(VI) reduction rate is not affected by NO_3_^−^ (Ceballos et al., 2020). Our results challenge this view. The reduction of Cr(VI) and NO_3_^−^ was concurrent, which was most evident at lower Cr concentration treatments (0.3 mM). The starting concentrations of Cr and NO_3_^−^ were 0.3 and 2.5 mM, respectively, and both were simultaneously lowered to negligible levels after 160 h. At higher Cr(VI) concentrations, the reduction rate of Cr(VI) remained stable at 0.12-0.14 mg*L^-1^*h^-1^, whereas denitrifiers failed to consume 2.5 mM NO_3_^−^ within 263 h. This is consistent with the competitive advantage caused by the increase in the concentration of one substrate under the substrate competition scenario (Straube, 2015).

Furthermore, we calculated the changes in redox potential (ΔE) and Gibbs free energy (ΔG) with pH under the biological standard condition and experimental condition (using 0.3 mM Cr treatment as an example) (Fig. S1 and detailed calculation in Supplementary Material). The results show that the redox potential and reaction energy yield of NO_3_^−^ and Cr(VI) are very similar under both biological standard conditions and experimental conditions. Specifically, under standard biological conditions, ΔE^0’^ _NO3− /NO2−_= +0.43v, ΔE^0’^_Cr(VI)/Cr(III)_ = +0.365v, ΔG^0’^ _NO3−/NO2−_ = -507.19 kJ/mol, ΔG^0’^_Cr(VI)/Cr(III)_ = -466,5 kJ/mol, and in the experimental condition, the redox potential and energy yield difference of Cr(VI) and NO_3_^−^ with CH_4_ is so small that can be ignored (ΔE difference<0.03v, ΔG difference<10kJ/mol). Our calculations demonstrate that the competition between NO_3_^−^ and Cr(VI) is highly sensitive to changes in pH and concentrations. In a neutral environment, neither Cr(VI) nor NO_3_^−^ has a clear thermodynamic advantage. Therefore, it is possible that the denitrifiers simultaneously utilize both as electron acceptors. This is also consistent with our experimental results, which show that the concentrations of NO_3_^−^ and Cr(VI) decrease at the same time, and the N-DAMO enrichment culture does not exhibit a clear preference for either electron acceptor. The calculation is also consistent with our experimental results, which show that the concentrations of NO_3_^−^ and Cr(VI) decrease at the same time, and the N-DAMO enrichment culture does not exhibit a clear preference for either electron acceptor.

There is another potential explanation for the simultaneous NO_3_^−^ and Cr(VI) reduction (Fig.6). Denitrification is a stepwise reaction in which NO2^−^ briefly exists in the system as an intermediate product, which is usually further reduced via the activity of nitrite reductase (Einsle & Kroneck, 2004). However, due to the strong oxidizing properties of Cr(VI), its presence may drive an abiotic NO_2_^−^ oxidation to NO_3_^−^ where Cr(VI) is reduced to Cr(III) (Eq.4; calculation in supplement material).

**Figure 6.**
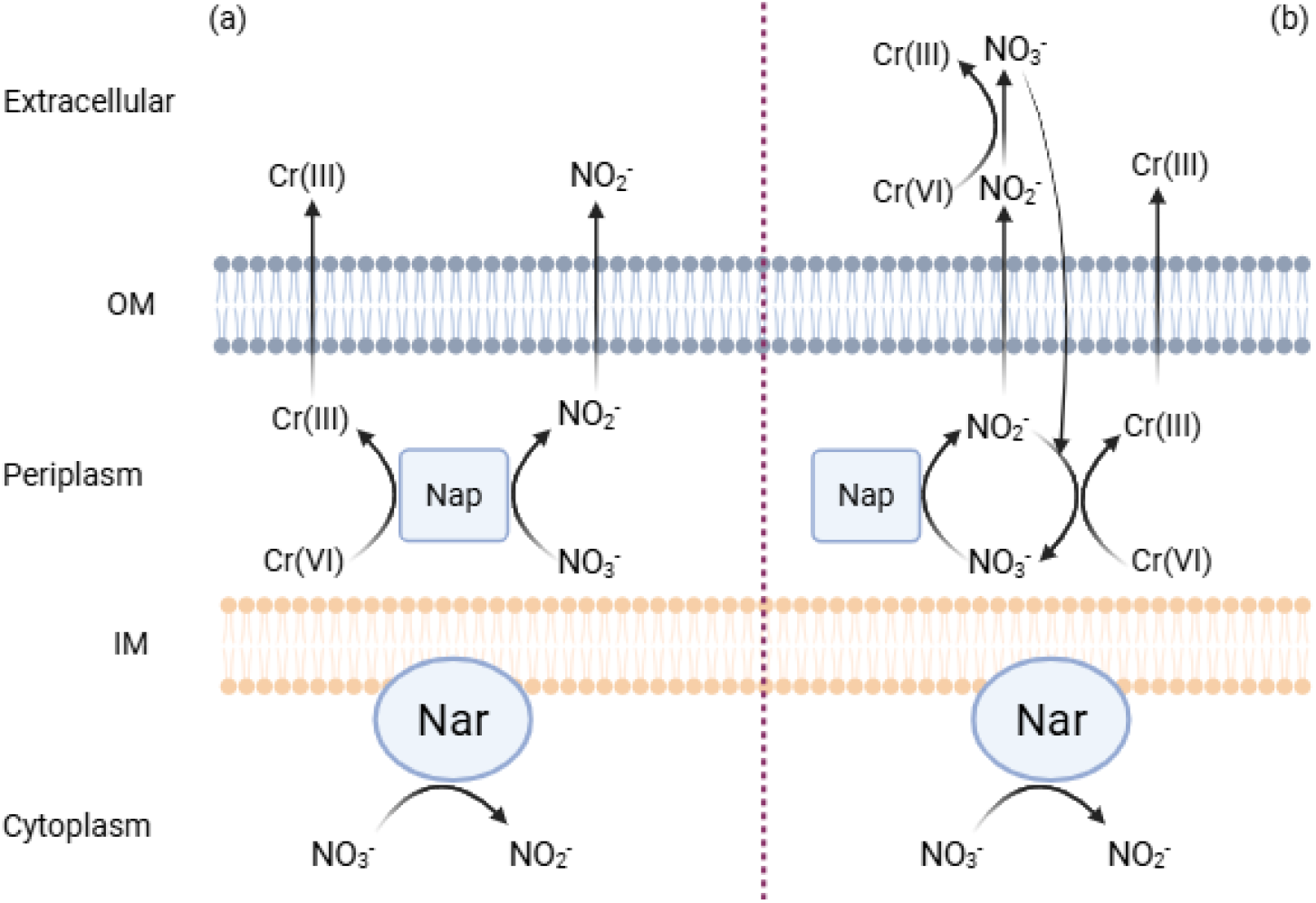
Two possible pathways for simultaneous Cr(VI) and NO_3_ ^-^ reduction by denitrifiers; a) NO_3_ ^-^ and Cr(VI) competing for the Nar; b) abiotic reduction of Cr(VI) by NO _2_^-^ in both the extracellular space and the periplasm. Most bacterial denitrifiers have both transmembrane nitrate reductases (Nar) and periplasmic nitrate reductases (Nap) (Lundberg et al., 2004). Nap is the dominant enzyme for nitrate reduction. The figure only illustrates the distribution of Nar and Nap in bacteria, while archaeal Nar is towards the extracellular space (Martinez-Espinosa et al., 2007).

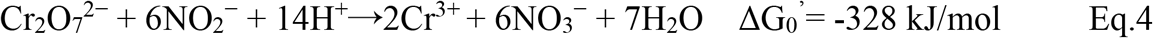

Future studies should aim to elucidate the mechanisms by which Cr(VI) inhibits denitrification, particularly by identifying the specific sites of Cr(VI) reduction (S. Wang et al., 2024). If Cr(III) is observed to form intracellular aggregates or precipitates in close proximity to denitrifying cells, this would suggest that Cr(VI) directly competes with NO_3_^−^ for nitrate reductase, thereby inhibiting the denitrification process. In contrast, if Cr(III) does not localize in specific cellular sites and instead forms soluble organo-Cr(III) complexes, for example with EPS, it would indicate that Cr(VI) may interfere abiotically with NO_2_^−^, potentially leading to its oxidation.

## 4. Conclusion

Heavy metals such as Cr(VI) often coexist with other environmental contaminants. To better understand the microbial-based simultaneous removal of common water pollutants such as CH_4_, NO_3_^−^ and Cr(VI), this study investigated the potential of the N-DAMO process to couple CH_4_ oxidation with Cr(VI) reduction. The results revealed that N-DAMO is highly sensitive to Cr(VI); even low concentrations significantly inhibited CH_4_ oxidation leading to a decline in key methanotrophic populations. In response, denitrifying bacteria within the flanking microbial community increased in relative abundance and became the primary drivers of both NO_3_^−^ and Cr(VI) reduction. Cr(VI) reduction likely occurred through two main pathways:

1. enzymatically, via nitrate reductase activity in denitrifying bacteria, resulting in competitive inhibition of denitrification; or
2. abiotically, through chemical reaction with NO_2_^−^, an intermediate product of denitrification.

These findings highlight important environmental and biotechnological implications. They suggest that while Cr(VI) can disrupt beneficial methane-oxidizing processes, denitrifying microbial communities offer resilience and functional redundancy that could be leveraged in engineered systems. Optimizing these microbial consortia may enhance the simultaneous bioremediation of nitrogen, carbon, and heavy metal pollutants in contaminated water and wastewater treatment applications.

## Acknowledgements

This study was supported by the SIAM Gravitation grant funded by NWO [Grant number 024.002.002] and an NWO-VIDI Talent grant [Grant number VI.Vidi.223.012].

## Supplement Material Calculation of the total amount of ^13^CO_2_ and ^15^N_2_O

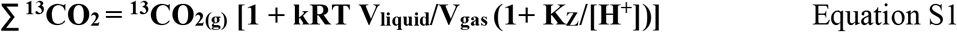

Where:

1. ∑^13^CO_2_ is the total amount of ^13^CO_2_ in the bottle
2. ^13^CO_2(g)_ is the amount of ^13^CO_2_ in the gas phase (headspace) in mmol.
3. k is the solubility coefficient of CO_2_, which is 3.3×10^−4^ mol/m^3^ Pa.
4. R is the universal gas constant, from the ideal gas law, which is 8.314 J mol^-1^ K^-1^.
5. T is the Kelvin temperature of incubation condition. 4ºC = 277.15 K, RT is ∼22 ºC = 295.15 K
6. V_liquid_ and V_gas_ are the volumes of the liquid and gas phases (in mL).
7. K_Z_ is the dissociation constant of the first step of carbonic acid dissociation. That is K_a1_, which you get from p*K*_*1*_, which you have to calculate (see below).
8. [H^+^] is the molar concentration of H^+^

### Gibbs Free energy for Cr(VI) reduction couple to NO_2_^−^ Oxidation

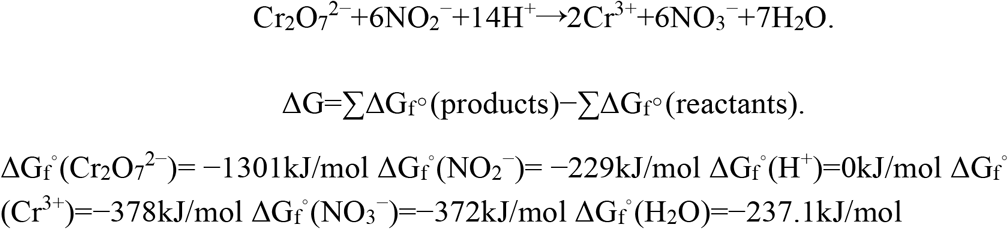

Data from Chemistry WebBook (https://webbook.nist.gov/) and Thermochemical Tables (https://atct.anl.gov/)

Net ΔG = ΔG_f_^°^(products)− ΔG_f_^°^(reactants)

=−4647.7−(−2675)

=−4647.7+2675

=−1972.7kJ/mol

328 kJ/mol per one mole of NO_3_^−^

ΔG is negative (ΔG=−1972.7 kJ/mol < 0), and the reaction is **spontaneous** under standard conditions.

### Calculation of redox potential for Cr(III)/Cr(VI) couple

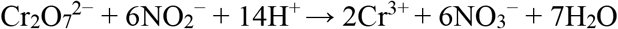

the electric potential of Cr_2_O_7_^2−^/Cr^3+^ is +1.33V (E^°^=+1.33V) (Ng et al., 2022) When pH **= 7**, the proton concentration is: [H^+^]=10^−7^M

The Nernst equation for this reaction is:

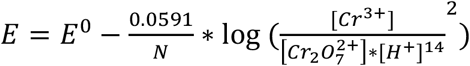

N= 6 (number of transferred electrons)

Assuming standard concentrations for ions (1M)

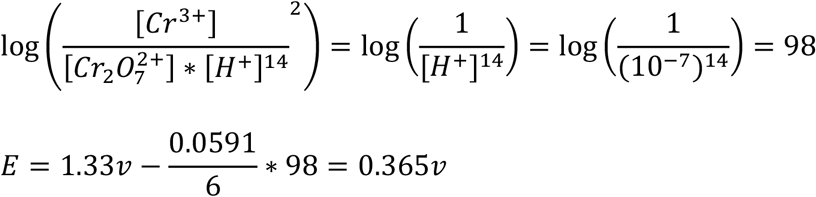

### Calculation details for Fig. S1

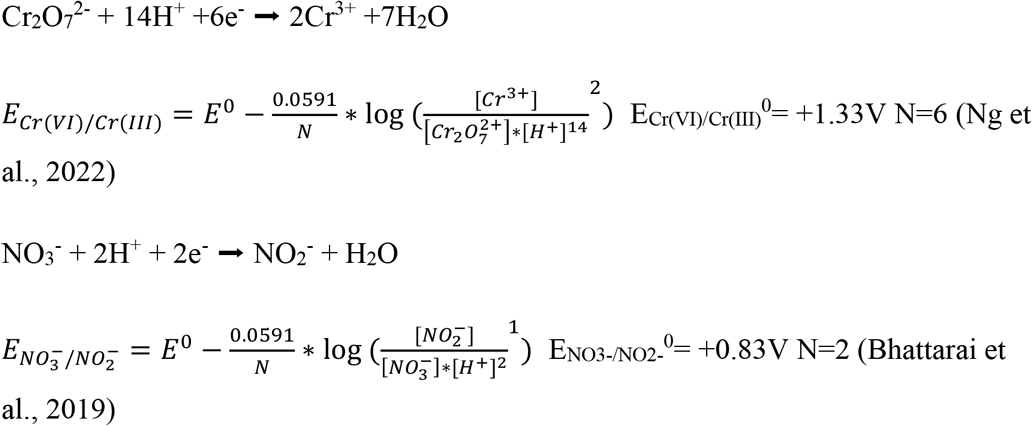

In standard biological conditions, [ox] = [red], therefore,

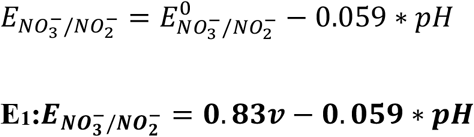

when pH=7, E_NO3−/NO2−_ = E^0’^ = 0.417V which is close to the previous calculation by Liebensteiner et al., 2014

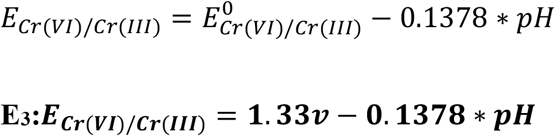

when pH=7, E_Cr(VI)/Cr(III)_ = E^0’^ = 0.365v

In experimental condition, here we took the concentration of 0.3mM treatment (26h) as an example, [NO_3_^−^]:[NO_2_^−^]=1.5:0.3, Cr(VI):Cr(III)=1000:1

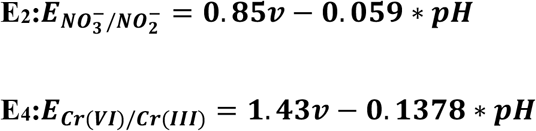

Similarly, according to ΔG =-nFE, we can then derive the relationship between Gibbs free energy and pH under various conditions. According to Eq.1 and Eq.2, n=8, ΔE_CH4_^0’^= -0.24v and F=96485C/mol, therefore:

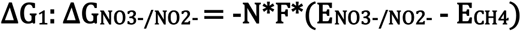

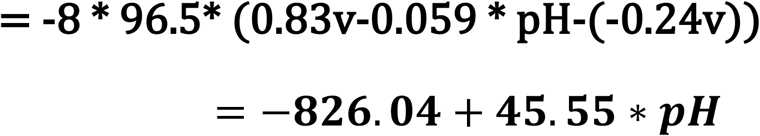

### Similarly

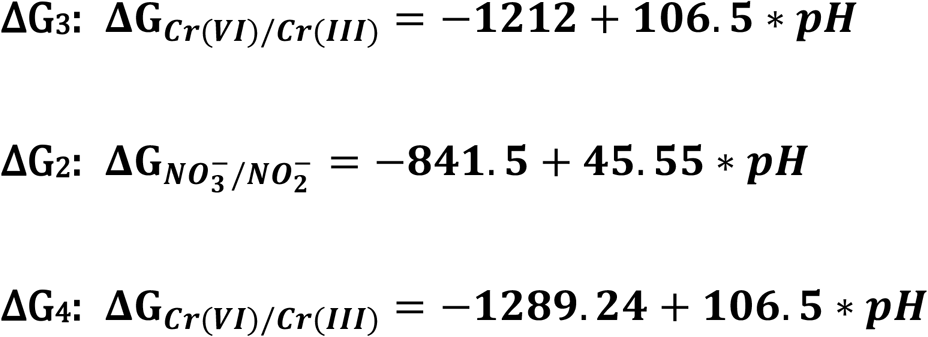

**Figure S1.**
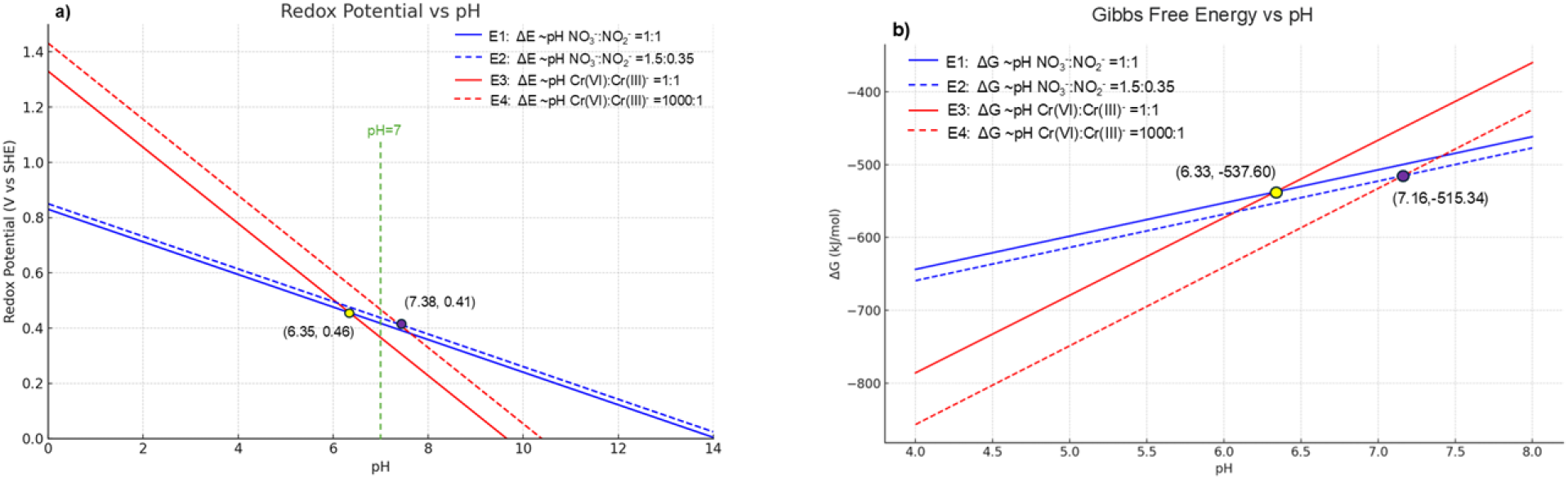
The change of a) redox potential and b) Gibbs free energy with pH in standard biological and experimental conditions. The red solid and dashed lines represent the Cr(VI)/Cr(III) couple at concentration ratios of 1:1 and 1000:1, respectively. The blue solid and dashed lines represent the NO_3_−/NO_2_− couple at concentration ratios of 1:1 and 1.5:0.35, respectively

